# TSPAN1 promotes human pancreatic cancer cells proliferation by modulating CDK1 via Akt

**DOI:** 10.1101/2020.05.29.123091

**Authors:** Xin Wang, Xiaozhuo Gao, Jiaxun Tian, Rui Zhang, Yun Qiao, Xiangdong Hua, Gang Shi

## Abstract

**Background:** To explore the potential therapeutic target to treat pancreatic cancers, Tspan1 was detected in human pancreatic cancer tissue and human pancreatic ductal adenocarcinoma cells and functional role of Tspan1 on proliferation was explored and the mechanism was investigated.

**Materials and Methods:** Tspan1 in PCC tissue and PDAC cell lines was measured by qRT-PCR and Western blot. Tspan1 was knock-downed and over-expressed in cells via transfection with Tspan1-siRNA and pLNCX-TSPAN1-cDNA, cell survival, proliferation and cell cycle were measured with MTT, Alamar blue and Flow Cytometry assay. The mRNA and protein expression were assessed by qRT-PCR and Western blotting. The expression of PI3K, Akt and p-Akt were detected, and CDK1 siRNA and specific inhibitor of Akt were used to explore the mechanism of TSPAN1 promoting PDAC cells proliferation.

**Results:** Tspan1 expression in PCC tissue and PDAC cells was increased. Transfection of siRNA targeting Tspan1 in BxPC3 and PNAC-1 cells obviously decreased cell proliferation and down-regulated CDK1 expression. Consistently, both cell proliferation and CDK1 expression in BxPC3 and PNAC-1 cells were up-regulated with pLNCX-TSPAN1-cDNA transfection. Cell cycle analysis showed that after knockdown of Tspan1 the G2/M phase ratio was increased to cause mitosis arrest, and TSPAN1 overexpression caused cell cycle transition from G2 to M phase to promote cell proliferation. And these were dependent on the modulation of CDK1 expression via Akt.

**Conclusion:** Tspan1 up-regulates CDK1 expression via activating Akt to promote human PCC cell proliferation and silencing of Tspan1 may be a potential therapeutic target to treat pancreatic cancers.

## 1 Introduction

Pancreatic cancer (PCC) is one of the most dangerous malignant tumor of the digestive system [1]. The American Cancer Society has reported that pancreatic cancer is becoming the third leading cause of cancer death in the United States, second only to lung cancer and colon cancer [2]. The annual survival rate of pancreatic cancer is only 8% and it is predicted that pancreatic cancer will be the second cause of cancer death by 2030 [1–2]. The clinical treatment of pancreatic cancer patients is mainly radical surgery supplemented by radiotherapy and chemotherapy [3]. Due to the difficulty in early detection of pancreatic cancer cells and the lack of effective screening indicators, at the time of diagnosis, the vast majority of patients are already at the advanced stage [2–3]. Therefore, only about 20% of pancreatic cancer patients have the opportunity of radical surgery [2–3]. For successful early diagnosis, the molecular biological events that contribute to the occurrence and development of PCC needs to be studied in detail to explore the ideal and effective diagnosis markers.

Tetraspanins, also referred to as the transmembrane 4 superfamily (TM4SF) proteins, are a family of proteins with four transmembrane domains [4,5]. Tetraspanins are often thought to act as scaffolding proteins, anchoring multiple proteins, such as various cell surface signaling molecules to one area of the cell membrane [4,6–7]. Lots of research work indicate Tspan1 is playing an important physiological role in cell adhesion, motility, activation, and proliferation, as well as contributing to pathological conditions such as metastasis or viral infection [9,10]. TSPAN1, a novel member of the TSPAN family [10], highly expressed in various cancers, such as gastric, colon, liver and esophageal cancers [8,11,12]. It was reported that Tspan1 plays important role in gastric and colon cancer cell invasion and metastasis [14,15]. Previous studies also revealed that TSPAN1 suppressed cell survival, proliferation, migration and invasion in colon cancer and skin carcinoma cells [14,15]. However, the role of TSPAN1 in PC cell proliferation is yet to be fully elucidated and detailed study is needed to explore its therapeutic potential for the treatment of pancreatic cancer.

In the present work, qRT-PCR and Western blot methods were applied to determine Tspan1 expression in the human pancreatic cancer tissues and the respective adjacent normal tissue, as well as in human pancreatic ductal adenocarcinoma (PDAC) cells. After transfection of Tspan1-siRNA and pLNCX-TSPAN1-cDNA plasmid into PDAC cell lines BxPC3 and PNAC-1, cell survival, cell proliferation and cell cycle were analyzed and the CDK1 expression were assessed. Furthermore, the expression levels of PI3K, Akt and p-Akt were investigated, and CDK1 siRNA and specific inhibitor of Akt were used to further explore the related mechanism of Tspan1 promotes human PDAC cells proliferation.

## 2 Materials and Methods

### Tissue collection, cell strains and cell culture

All cell strains were obtained from the Shanghai Institute of Biochemistry and Cell Biology (Shanghai, People’s Republic of China), including human pancreatic ductal adenocarcinoma (PDAC) strains BxPC3 and PNAC-1cells, HEK293 cells and normal human pancreatic cell line HPC-Y5. Culture medium was Dulbecco’s modified Eagle medium (DMEM, Invitrogen; Thermo Fisher Scientific, Inc., Waltham, MA, USA) supplemented with 10% FCS (FCS; Invitrogen; Thermo Fisher Scientific, Inc.). When the cultured cells reached 70-80% confluence at 37° C in a humidified 5 %CO2 incubator, the cells were passaged with 0.25% (w / v) trypsin of 0.02% (w / v) EDTA solution. For analysis of the role of Akt on Tspan1 modulating CDK1 expression, 1 h after passage of the transfected cells, AKt inhibitor (MK-2206 2HCl; 10 μM; Cat.no. S1078, Precision Technologies Pte Ltd, Houston, TX, USA) was added to the culture medium and the cells were cultured at 37°C for a further 24 h. The protein expression of CDK1 was detected by Western blot. Thirty pairs of human pancreatic carcinoma (PCC) tissue specimen and their corresponding normal tissue specimen were collected during routine surgery at Liaoning Cancer Hospital and China Medical University. Each patient signed a official informed consent form prior to surgery. They were enrolled between July 2017 and July 2018 (16 men and 14 women; aged 45-75 years, mean 51 years), all diagnosed with pancreatic carcinoma and had no radiotherapy or chemotherapy treatment. The ethics committee (EC) of China Medical University approved the study and this study was conducted within the ethical guidelines of EC, China Medical University.

### RNA isolation and Quantitative reverse-transcription polymerase chain reaction (qRT-PCR)

RNA was isolated using the RNeasy mini kits (Qiagen GmbH, Hilden, Germany) according to themanufacturer’s experimental manual and then subjected to reverse transcription according to themanufacturer’s instruction. Real time PCR was carried out on the Applied Biosystems 7900HT thermal cycler instrument (Applied Biosystems, USA) according to the manufacturer’s instructions. Briefly, qRT-PCR was executed in a total of 20μl reaction volume which includes 2 X Mix SYBR green I (10μl; Promega Corporation, Madison, WI, USA), primer (0.25μl, 10pmol/l), template DNA (1 μl) and autoclaved Millipore water. The realtime PCR reaction conditions were denaturation at 95 °C for 2 minutes, then 95 °C, 15 seconds for a total of 50 cycles, finally 60 °C for 45 seconds. The expression of the target gene is evaluated by this formula to calculate the fold change: 2-[[Ct (control) gene X-Ct (control) gapdh][Ct (activated) gene X-Ct (activated) gapdh]] [16]. The sequences of used primers were listed as following: Human *Tspan1* forward, 5’-CGTTGTGGTCTTTGCTCTTG-3 ‘and reverse, 5’-TTCTTGATGGCAGGCACTAC-3’; Human *CDK1* forward,5’ TTTTCAGAGCTTTGGGCACT −3’ and reverse: 5’-CCATTTTGCCAGAAATTCGT-3’. Human *GAPDH* (glyceraldehyde-3-phosphate dehydrogenase) forward: 5’-GAAGGTGAAGGTCGGAGTC-3’ and reverse, 5’-GAAGATGGTGATGGGATTTC-3’. The mRNA expression of *Tspan1* and *CDK1* was normalized to the expression of endogenous *GAPDH* mRNA to obtain their relative expression.

### Western blotting

RIPA buffer and protease inhibitor (Sigma-Aldrich; Merck KGaA, Darmstadt, Germany) were applied to lyse the cells then the proteins were collected. Eighty micrograms of protein from each group was denatured at 100 °C for 5 minutes and revolved with a 10% SDS-PAGE stacking gel (Millipore Corporation, Billerica MA) and then transferred to a PVDF membrane (Millipore Corporation,Billerica MA) by electroblotting method. The PVDF membrane was incubated with 5% fresh skim milk at room temperature for 1 hour to block non-specific binding, followed by addition to 1st antibodies directed against Tspan1 (1:300; cat.no.NBP2-33867; Novus Biologicals, LLC, Littleton, CO, USA), CDK1 (1:300; cat.no.PA5-82086; ThermoFisher Scientific, Waltham, MA USA), PI3K (1:500; cat. no.4292; Cell Signaling Technology, Inc., Danvers, MA, USA), phosphorylated (p)-Akt (1:500; cat. no.4060S; Cell Signaling Technology, Inc.), Akt (1:500; cat. no. 4691s; Cell SignalingTechnology, Inc.) and anti-beta actin antibody (1:1000, cat. no. NB600-503, Novus Biologicals, LLC) was respectively incubated overnight at 4 degree. TBST buffer was used to wash away non-specific binding, then the PVDF membrane was incubated together with HRP-conjugated secondary antibody (HRP-linked anti-rabbit IgG antibody, 1:3000, cat. no. NB710-57836; Novus Biologicals, LLC) at room temperature for 1.5 hours. ECL system (Enhanced Chemiluminescence, Pierce;Thermo Fisher Scientific, Inc.) was applied to develop protein signals. Each band of protein was scanned then quantified the optical density using a scanning densitometer and Quantity One Software, version 4.4.1 (Bio-Rad Laboratories, Inc., Hercules, CA, USA). The relative expression level of each target protein was obtained by normalizing with the corresponding beta-actin density.

### Overexpression of TSPAN1 in BxPC3 cells and PNAC-1 cells

The TSPAN1 cDNA cloning plasmid (cat. no. HG 13073-UT) was purchased from Beijing Zeping Bioscience & Technology Co., Ltd (Beijing, China). A recombinant plasmid (pLNCX-TSPAN1-cDNA) was constructed and sequenced at BioSune Biotechnologies (Shanghai, China). Ten μg of the pLNCX-TSPAN1-cDNA recombinant plasmid or control plasmid pLNCX was transfected into BxPC3 and PNAC-1 cells by calcium phosphate precipitation method [17]. Specifically, BxPC3 and PNAC-1 cells were cultured in a 35 mm cell culture dish to 70-80% confluence. DNA-calcium phosphate transfection complex was prepared by gently adding plasmid DNA and 2M CaCl2 (Sigma-Aldrich; Merck KGaA) dropwise to HBS buffer (Sigma-Aldrich; Merck KGaA). This transfection complex was added dropwise to 70-80% confluent cells then the cells were incubated at 37°C for 8 hours, then switched to fresh DMEM medium for culture. pLNCX plasmid was transfected into cells to serve as the control. The cells were subjected to mRNA expression analysis 48 hours after transfection, and protein expression analysis was performed 72 hours after transfection.

### Knockdown of Tspan1 in BxPC3 cells and PNAC-1 cells by RNAi

Down-regulation of Tspan1 in BxPC3 cells and PNAC-1 cells was achieved using specific siRNAs targeting Tspan1 (Tspan1 siRNA, cat.no.4392420; Thermo Fisher Scientific, Inc.). The scramble-siRNA (siNC, cat.no.4390846; Thermo Fisher Scientific, Inc.) transfected cells were used as controls. siRNA was transfected into BxPC3 cells and PNAC-1 cells following the manufacturer’s instructions. Firstly, 1 × 10^5^ cells/ml of BxPC3 cells and PNAC-1 cells were resuspended in DMEM cell culture medium. RNAi MAX Lipofectamine transfection reagent (Ambion; Thermo Fisher Scientific, Inc.), siRNA (20 nM) and DMEM cell culture medium were mixed together to prepare a transfection complex. The transfection complex and BxPC3 cells and PNAC-1 cells were then mixed together and seeded in 6-well (2 x 10^5^ cells/well) cell culture plates (Nunc, Rochester, NY, USA) and cultured at 37°C for 24 hours. After 48 hours, RNA was extracted to detect mRNA expression, and after 72 hours, protein was extracted to detect protein expression. The qRT-PCR method was used to confirm the silencing efficiency. Both the transfection method of CDK1 siRNA (cat.no. AM16708; Thermo Fisher Scientific, Inc.) into HEK293, Tspan1-overexpressed BxPC3 cells and PNAC-1 cells, and the detection of silencing efficacy were the same as aforementioned.

### Cell proliferation (MTT) assay

The effects of up-regulation and down-regulation of Tspan1 on cell proliferation and viability in BxPC3 cells and PNAC-1 cells were determined by MTT assay. MTT assays were performed at 0, 24, 48, 72, 96 and 120 hours intervals after transfection. The specific experimental method was as follows: 3×10^4^ cells were inoculated into a 96-well plate one day before transfection, then 150μl DMEM containing 20μl MTT (5mg/ml; Sigma-Aldrich; Merck KGaA) was added in per well and the cells were cultured in a 5% CO2 incubator at 37°C for 4 hours. The medium was then gently aspirated, and 150μl of dimethyl sulfoxide (DMSO, Sigma-Aldrich; Merck KGaA) was added to each well to dissolve the crystals of the crystal. The absorbance of formazan amount from 490 nm wavelength was measured using a Bio-Rad Microplate Reader 550 (Bio-Rad Laboratories Inc. Tokyo, Japan). Cell growth curves were plotted using mean ± standard deviation data.

### Alamar blue cell survival assay

Alamar blue assay was performed to assess cell viability and proliferation. The Alamar blue assay measures quantitatively cell proliferation as well as relative cytotoxicity. It incorporates a water soluble colorimetric oxidation-reduction (Redox) indicator that changes color in response to the chemical reduction of the culture medium resulting from cell growth (metabolic activity). Briefly, alamar blue dye (Invitrogen, USA) at a concentration of 10% v/v in PBS was added to each well seeded with control cells or transfected cells and incubated for 4 hours at 37°C in 5% CO_2_. 100μl of the medium was transferred to a fresh 96 well plate and absorbance was read at 570nm (reduction) and 600nm (oxidation) using a spectrophotometer (SpectraMax M5, Molecular Devices, LLC, CA, United States). The data were expressed as a fraction of the cell activity.

### Flow cytometry analysis of cell cycle

Control cells and transfected cells were washed twice with ice-cold PBS then fixed with 70 % ethanol at −20°C for 24 hours. Cells were further washed twice with icecold PBS then stained with propidium iodide (PI, 20μg/ml) added in RNAase (100 μg/ml) in darkness for 30 minutes at room temperature. Cell cycle phase analysis was performed using a BD FACSCalibur flow cytometer (Becton & Dickinson, USA). 20000 cells per sample were collected and analyzed by using the CellQuest software (version 5.1; BD Biosciences) analysis program.

### Statistical Analysis

The results are expressed as the mean ± standard deviation (n=3). At least three independent experiments were performed in each experiment. Statistical differences between the two groups were analyzed using SPSS 17.0 software (SPSS, Inc., Chicago, IL, USA) and calculated using a two-tailed Student’s t-test. One-way variance analysis, Tukey’s post hoc tests and SPSS 17.0 software (SPSS, Inc.) were applied to analyze the significant differences among groups. P < 0.05 was considered to be statistically different.

## 3 Results

### Expression of Tspan1 in clinical pancreatic cancer (PCC) tissue and human pancreatic ductal adenocarcinoma (PDAC) cells

The mRNA expression levels of Tspan1, measured by qRT-PCR, were significantly increased in the human pancreatic cancer tissue (Fig.1A) compared to its adjacent normal pancreatic tissue. The mRNA and protein expression levels of Tspan1 were measured in human pancreatic ductal adenocarcinoma (PDAC) cells lines BxPC3 and PNAC-1cells and normal human pancreatic cell line HPC-Y5 using qRT-PCR and Western blot methods. The results showed that BxPC3 and PNAC-1 cells expressed higher mRNA and protein levels of Tspan1 compared to HPC-Y5 cells (Fig.1B and 1C).

**Figure 1.**
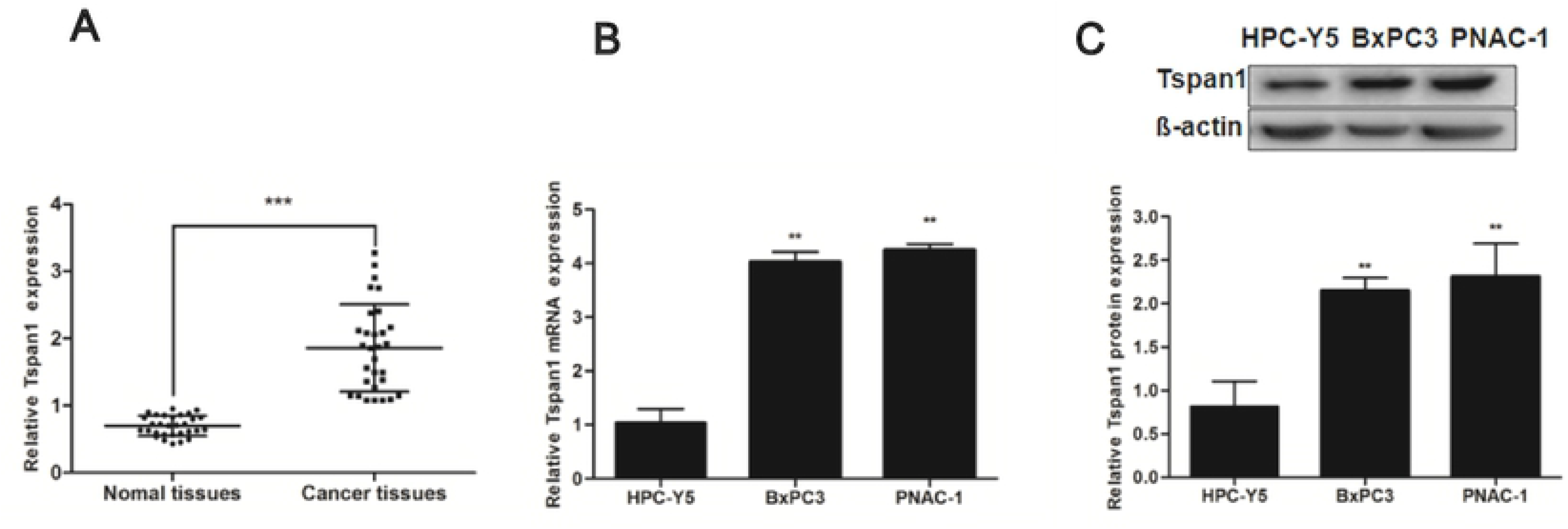
Expression level of Tspan1 in human pancreatic cancer tissue specimen and PDAC cells lines. The expression level of Tspan1 was detected. A. High mRNA expression of Tspan1 was observed in human pancreatic cancer tissue specimen. B. The mRNA expression level of Tspan1 was assessed by qRT-PCR assay in human pancreatic ductal adenocarcinoma (PDAC) cells lines BxPC3 and PNAC-1cells, and normal human pancreatic cell line HPC-Y5. C. Western blot result of Tspan1 expression. Data are reported as the mean ± standard deviation. *P<0.05 (n=3) vs. HPC-Y5,**P<0.01 (n=3) vs. HPC-Y5.

### Overexpression of Tspan1 in BxPC3 cells and PNAC-1 cells

Calcium phosphate precipitation method was used to transfect the pLNCX-TSPAN1-cDNA recombinant plasmid and control pLNCX plasmid into BxPC3 and PNAC-1 cells. After transfection, the expression of Tspan1 was determined with qRT-PCR (Fig 2A) and Western blot (Fig 2B). Cells transfected with control pLNCX plasmid served as control. The expression of Tspan1 was obviously up-regulated in pLNCX-TSPAN1-cDNA transfected cells (Tspan1) compared with control pLNCX plasmid transfected cells (Ctrl).

**Figure 2.**
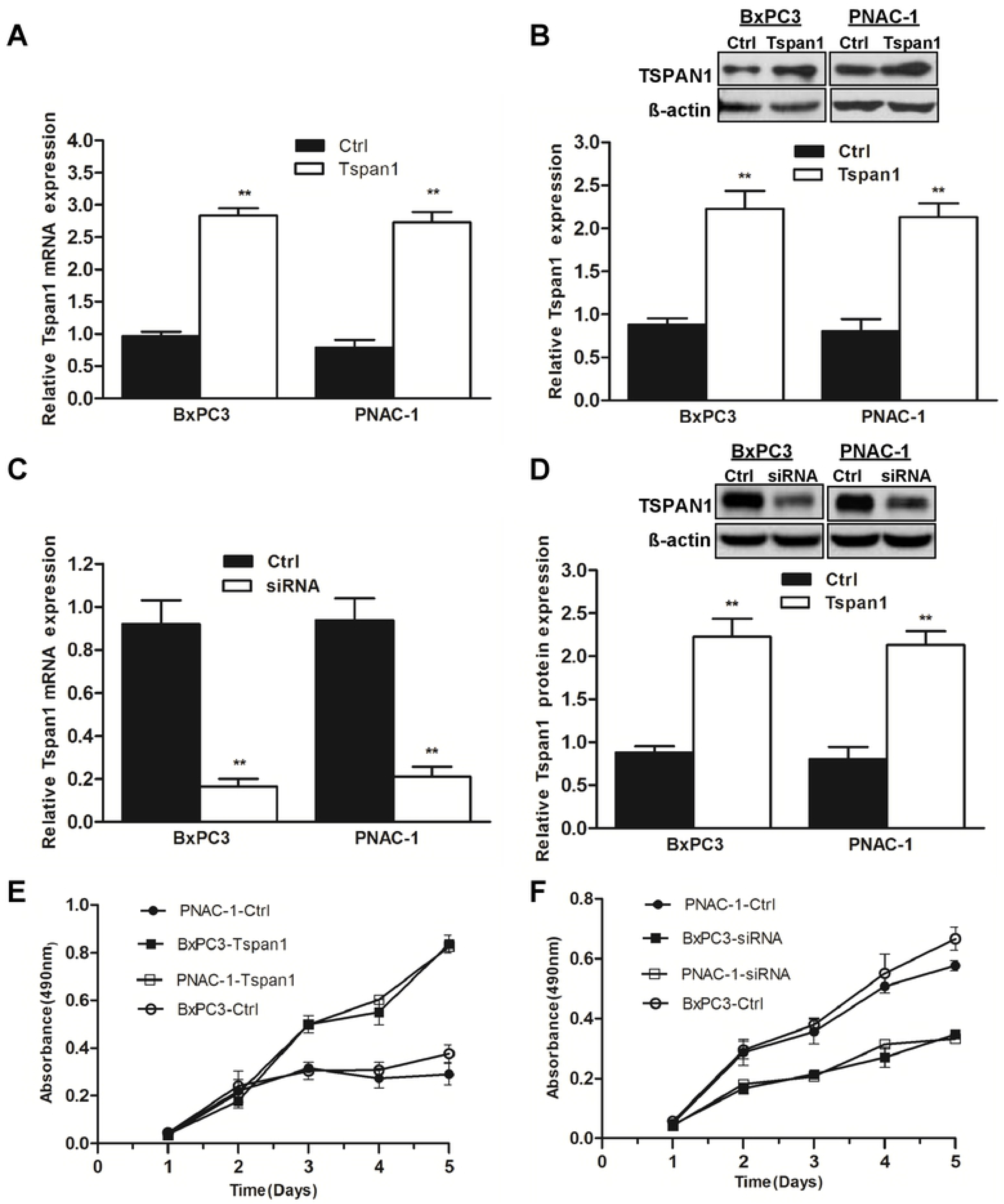
Tspan1 modulated BxPC3 cells and PNAC-1 cells proliferation. A. pLNCX-TSPAN1-cDNA recombinant plasmid and control pLNCX plasmid were transfected into BxPC3 and PNAC-1cells. The mRNA levels of Tspan1 were analyzed by qRT-PCR analysis. Control pLNCX plasmid-transfected BxPC3 and PNAC-1 cells serve as control cells (Ctrl). After pLNCX-TSPAN1-cDNA was transfected into BxPC3 and PNAC-1 cells (Tspan1), Tspan1 mRNA expression was up-regulated. B. Western blot results showed that Tspan1 protein expression was increased after pLNCX-TSPAN1-cDNA transfection. C. The siRNA targeted to Tspan1 was transfected into BxPC3 and PNAC-1 cells (siRNA). BxPC3 and PNAC-1 cells transfected with scramble siRNA were treated as control (Ctrl). Quantitative RT-PCR method was used to analyze the mRNA expression levels of Tspan1. D. Western blot results showed that Tspan1 protein expression was decreased after Tspan1-siRNA transfection. E. Cell proliferation was measured by MTT method after TSPAN1 overexpression, BxPC3 and PNAC-1 cells proliferation was significantly increased after Tspan1 overexpression. F. Cell proliferation assay after knockdown of Tspan1. BxPC3 and PNAC-1 cells proliferation was significantly suppressed by Tsapn1-siRNA transfection. *P<0.05 (n=3) vs. Ctrl, **P<0.01 (n=3) vs. Ctrl.

### Silencing of Tspan1 expression by transfection of Tspan1 siRNA

Tspan1 siRNA was transfected into the human PDAC cell lines BxPC3 and PNAC-1. Scramble-siRNA was used to confirm the specificity of Tspan1-siRNA. The silencing efficiency of Tspan1-siRNA transfection was confirmed via qRT-PCR (Fig. 2C). Tspan1 mRNA was significantly decreased in the Tspan1-siRNAs transfected cells compared to the scramble-siRNA-transfected control cells (Ctrl). Western blot analysis also showed that the expression of Tspan1 was significantly decreased after transfection of Tspan1-siRNA in BxPC3 and PNAC-1 cells (Fig. 2D).

### Tspan1 promoted BxPC3 and PNAC-1 cell proliferation

The effects of Tspan1 overexpression and Tspan1 silencing on the BxPC3 and PNAC-1 cells proliferation were measured using MTT assay. The cell growth curves results showed that overexpression of Tspan1 obviously promoted cell proliferation of BxPC3 and PNAC-1cells (Fig. 2E). Meanwhile, Tspan1 silencing significantly suppressed cell proliferation of BxPC3 and PNAC-1 cells (Fig. 2F).

### TSPAN1 overexpression caused evident cell cycle transition from G2 to M phase

To further investigate the Tspan1 promoting cell proliferation, cell cycle distribution was analyzed by flow cytometry. in Figure 3A, flow cytometry analysis showed that scramble-siRNA-transfected BxPC3 cells and PNAC-1 cells (Ctrl) were presented in G2/M (9.73±1.08 and 11.11±1.79%) phase. In Tspan1-siRNA-transfected BxPC3 cells and PNAC-1 cells (siRNA), the G2/M ratio was 17.69±1.84% and 19.85±1.53%, respectively (p< 0.05). The proportion of cells in G2/M phase significantly increased. It indicated that the fractions of cells in M-phase of mitosis - which represents a population of dividing cells - had significantly decreased after knockdown of Tspan1. And the cell cycle was maintained in a longer G2 phase for intranuclear replication, which arrested cells growth in the mitosis phase and inhibited cell proliferation. While TSPAN1 overexpression caused evident cell cycle transition from G2 to M phase to promote cell proliferation (Fig. 3B). In detail, In the pLNCX plasmid transfected BxPC3 and PNAC-1 cells (Ctrl), the proportion of G2/M phase cells was 12.99±1.27% and 10.58±1.12%. In the pLNCX-TSPAN1-cDNA transfected BxPC3 and PNAC-1 cells (Tspan1), the proportion of G2/M phase cells was 2.05±0.133% and 1.32±0.127%. The data showed that after overexpression of Tspan1, the proportion of cells in G2 phase decreased, and the proportion of cells in M phase significantly increased, it suggests that Tspan1 overexpression promoted cell cycle transition from G2 to M phase to increase cell proliferation.

**Figure 3.**
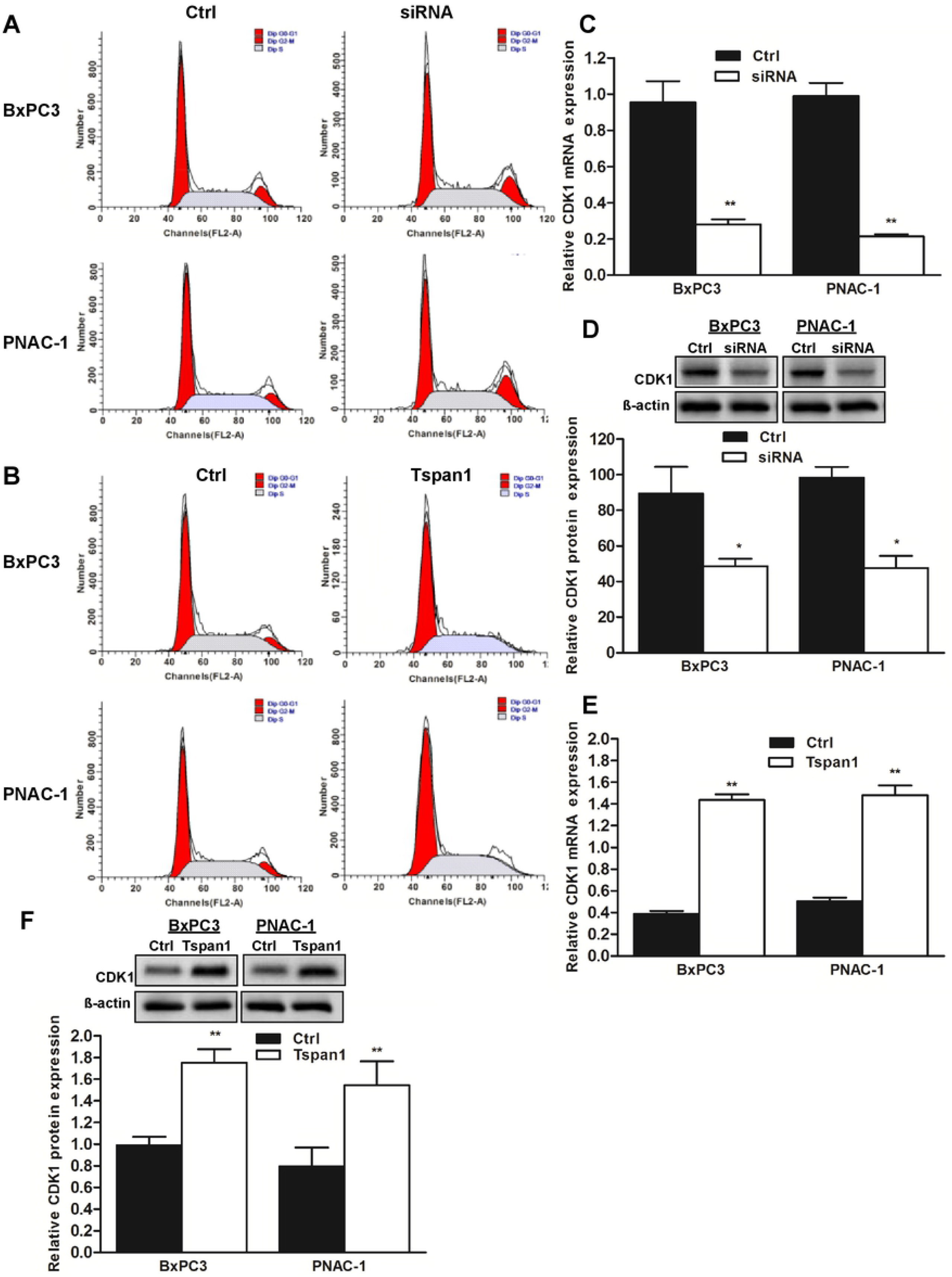
Tspan1 regulated cell cycle progession and CDK1 expression. BxPC3 and PNAC-1 cells were transfected with Tspan1 siRNA (siRNA) and scramble siRNA (Ctrl), pLNCX-TSPAN1-cDNA recombinant plasmid (Tspan1) and pLNCX plasmid (Ctrl). The cell cycle was analyzed by flow cytometry. A. Cell cycle assay after knockdown of Tspan1. B. Cell cycle assay after overexpression of Tspan1. BxPC3 and PNAC-1 cells were transfected with Tspan1 siRNAs (siRNA) and scrambled siRNA (Ctrl). The mRNA expression (C) and protein expression (D) of CDK1 after silencing of Tspan1 in BxPC3 and PNAC-1 cells were quantified, respectively. After pLNCX-TSPAN1-cDNA transfection, the mRNA levels of CDK1 were detected using qRT-PCR (E). The protein levels of Tspan1 were detected and quantity analysis of relative protein levels of Tspan1 was conducted (F). *P< 0.05 vs Ctrl; **P< 0.01 vs Ctrl.

### Tspan1 modulates the expression of CDK1 to regulate cell proliferation

To further explore Tspan1 regulating the progression of cell cycle, the expression level of CDK1 was measured after Tspan1 overexpression and Tspan1 silencing. Tspan1 ectopic overexpression in the BxPC3 cells and PNAC-1 cells increased the mRNA and protein expression of CDK1 (Fig. 3E and 3F). As expected, the mRNA and protein expression of CDK1 were significantly decreased in Tspan1-siRNA-transfected BxPC3 cells and PNAC-1 cells in comparison to scramble-siRNA-transfected control cells (Ctrl)(Fig. 3C and 3D). It indicates that Tspan1 regulates BxPC3 cells and PNAC-1 cells cycle progression by modulating cell cycle-associated protein CDK1. For confirm our hypothesis, we knocked down the expression of CDK1 in the Tspan1-overexpressed BxPC3 cells and PNAC-1 cells using CDK1-siRNA transfection, and Alamar blue assay method was used to observe the cell proliferation and survival. The results showed that the up-regulated cell survival and proliferation, induced by overexpression of Tspan1 in BxPC3 cells and PNAC-1 cells, was significantly attenuated by the transfection of CDK1 siRNA (Figure 4C and 4D). The data indicates that overexpression of Tspan1 which promotes cell proliferation dependents on the up-regulation of CDK1 expression.

**Figure 4.**
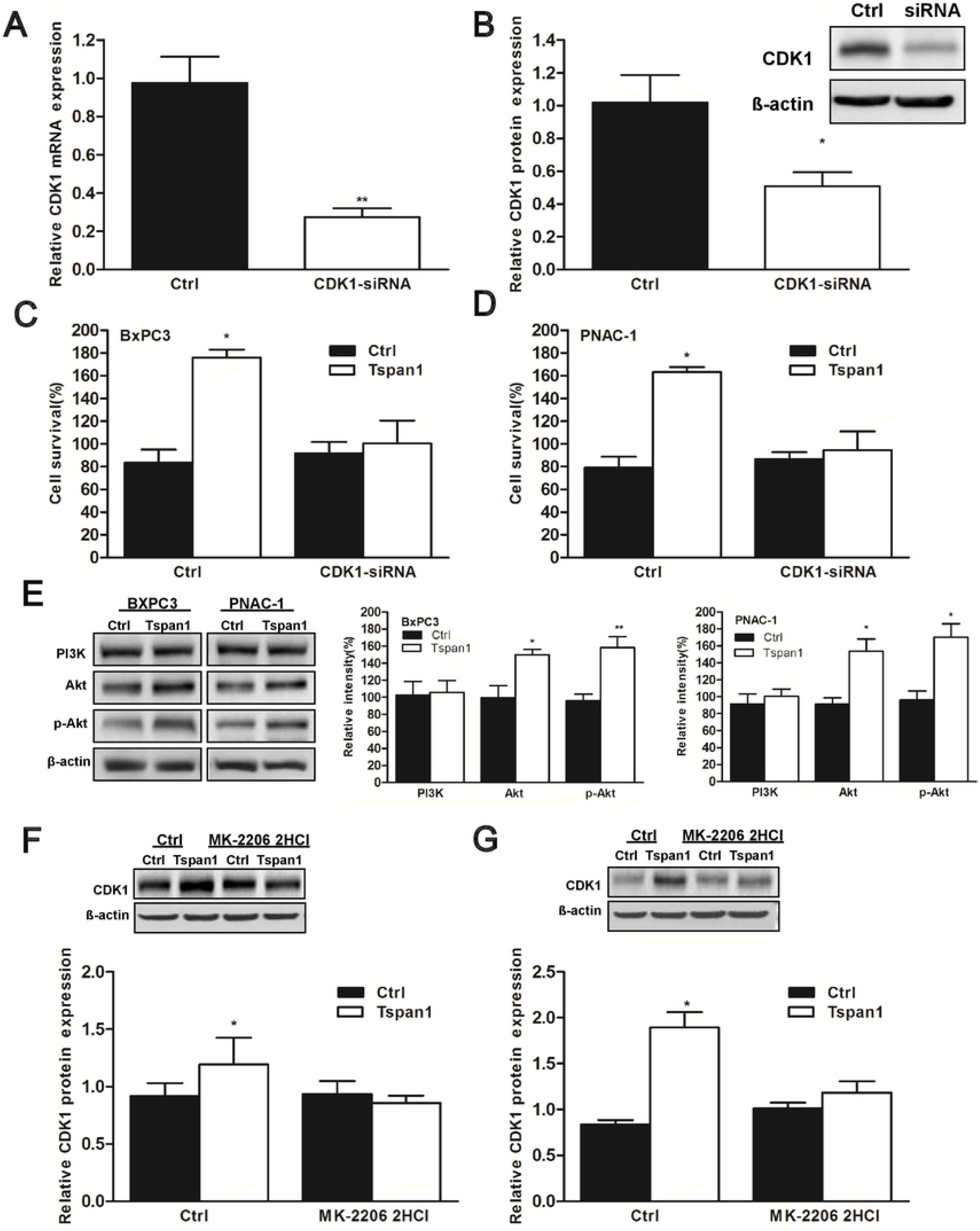
Tspan1 up-regulated BxPC3 and PNAC-1 cells proliferation and CDK1 expression depend on the activation of Akt. HEK293 cells were transfected with CDK1 siRNA, the expression levels of CDK1 were detected with qRT-PCR (A) and Western blot (B). BxPC3 and PNAC-1 cells were transfected with pLNCX-TSPAN1-cDNA plasmid (Tspan1). Plasmid pLNCX was transfected into BxPC3 and PNAC-1 cells as control cells (Ctrl). After transfection of CDK1 siRNA, cell proliferation ability and CDK1 expression were observed and analyzed. Alamar blue assay method results of cell survival and proliferation in the Tspan1-overexpressed BxPC3 cells (C) and PNAC-1 cells (D) after CDK1-silencing. CDK1 knock-down significantly abolished the increased cell survival and proliferation ability induced by Tspan1 overexpression. E. The expression levels of PI3K, Akt and p-Akt in the Tspan1-overexpressed BxPC3 cells and PNAC-1 cells were measured by western blotting. It demonstrated that Tspan1 overexpression could promote the expression and phosphorylation of Akt. Then MK-2206 2HCl, a specific inhibitor of Akt were used to suppress the Akt activation, then the expression of CDK1 was assessed in the Tspan1-overexpressed BxPC3 cells (F) and PNAC-1 cells (G). It showed that suppression of Akt significantly attenuated the increased CDK1 expression induced by Tspan1 over-expression. * P< 0.05 (n=3) vs Ctrl, **P<0.01 (n = 3) vs Ctrl.

### Tspan1 promotes BxPC3 and PNAC-1 cells proliferation and CDK1 expression via activation of Akt

To further investigate the mechanisms of Tspan1 modulate CDK1 expression and cell cycle, we investigated the expression levels of PI3K, Akt and p-Akt in the Tspan1-overexpressed BxPC3 cells and PNAC-1 cells by western blotting. The results (Fig. 4E) showed that the overexpression of Tspan1 could increase the content of Akt and p-Akt. It demonstrated that Tspan1 overexpression could promote the expression and phosphorylation of Akt, thus activate Akt in the BxPC3 cells and PNAC-1 cells. Therefore, the aforementioned results implied that Tspan1 may modulate the CDK1 expression through the activation of Akt. Furthermore, MK-2206 2HCl, a specific inhibitor of Akt were used to suppress the Akt activation, then the expression of CDK1 was assessed. It showed that suppression of Akt significantly attenuated the increased CDK1 expression induced by Tspan1 overexpression (Fig. 4F and 4G). In summary, these results indicate that Tspan1 modulated the expression of CDK1 via the activation of Akt.

## 4 Discussion

Pancreatic cancer is a highly malignant tumor of the digestive system with a very poor prognosis. About 90% of it originates from the glandular epithelium [3]. It has been reported that pancreatic cancer has now surpassed breast cancer as the third leading cause of cancer death in the United States [1–2]. Due to the low early diagnosis rate of pancreatic cancer, most the patients have lost the opportunity to have surgery at the time of diagnosis, and various existing treatment strategies, such as radiotherapy, chemotherapy, targeted therapy, etc., have not significantly improved their survival rate [3]. The early diagnosis of pancreatic cancer can significantly improve the prognosis [18,19]. Therefore, exploring the ideal molecular markers for effective early diagnose has become a research hot spot in recent years.

TSPANs are a membrane protein superfamily that has four transmembrane domains [20]. This four-transmembrane protein is a type of transmembrane glycoprotein that is widely expressed on the cell surface and is characterized by a four-segment transmembrane structure, including two extracellular hydrophilic rings (EC1 and EC2) and an intracellular N-terminus and C-terminus [21]. TSPANs play important role in various biological activities such as composing cell structure, regulating cell movement and proliferation, and regulating immune response [4]. Tspan1 is a novel member of the TSPANs superfamily [10]. Previous studies have reported that Tspan1 was high-expressed in various types of cancers, such as hepatocellular carcinoma, colon cancer, gastric carcinoma and skin squamous carcinoma [13–16, 22]. In our study, the results indicated that Tspan1 was also highly expressed in human pancreatic cancer tissue compared with the adjacent normal pancreatic tissue. And Tspan1 was also high-expressed in the human PDAC cell lines BxPC3 and PANC-1 compared with that in normal pancreatic cell line HPC-Y5. These founding may indicate that the expression of Tspan1 could be used as a specific marker for human PCC diagnosis and treatment. For confirming this hypothesis, we detected the effects of Tspan1 over-expression and Tspan1-silencing on the cell survival and proliferation in PDAC cells BxPC3 and PNAC-1. Our results demonstrated that the ectopic expression of Tspan1 promoted PDAC cell survival and proliferation. While the silencing of Tspan1 reduced PDAC cell survival and proliferation, which was consistent with previous findings in different cancer cells [15,16,23]. Therefore, it implied that Tspan1 may act as an oncogenic gene. To explore the underlying mechanisms that Tspan1 promoting PDAC cells proliferation, cell cycle was detected and analyzed after Tspan1 over-expression and Tspan1-silencing. Our results showed that Tspan1 over-expression caused evident cell cycle transition from G2 to M phase to promote cell proliferation. While Tspan1-silencing inhibits cell proliferation by arresting the cell cycle in the G2 phase. Therefore, the modulation of Tspan1 expression could regulate cell cycle and affect cell proliferation, which also confirmed that Tspan1 plays an important role in pancreatic cancer development.

The key factors in the regulation of cell cycle are cyclin-dependent kinases (CDKs) and cyclins [24]. CDK1 is one of the most critical CDKs for cell cycle regulation [25,26]. In the late G2 phase and early M phase, cyclin binds to CDK1 then initiates cell progression to M phase, which controls cell cycle arrest and closure. Deregulation of CDK1 directly leads to genomic instability and uncontrolled cell proliferation [25–27]. Some researchers have reported that the abnormal expression of cyclinB/CDK1 promotes cell proliferation [27,28]. Our results show that the transfection of Tspan1-siRNA obviously down-regulated CDK1 expression and inhibited cell proliferation in BxPC3 and PNAC-1 cells, while Tspan1-overexpression increased CDK1 expression, caused evident cell cycle transition from G2 to M phase to promote cell proliferation. It indicates that Tspan1 promotes BxPC3 cells and PNAC-1 cells proliferation and cell cycle progression by modulating the expression of CDK1. Using CDK1-siRNA, our results show that CDK1-silencing significantly attenuated the cell proliferation upregulated by Tspan1-overexpression in the BxPC3 and PNAC-1 cells. It furthermore confirms that overexpression of Tspan1 promoting cell proliferation was dependent on the up-regulation of CDK1.

Walker MP [34] and Yani Tang [30] reported that the PI3K/Akt signaling pathway could activate the downstream transcription factors, nuclear factor-κB and Sip1/tuftelin-interacting protein (STIP) to regulate the expression of CDK. In our study, for exploring the role of the PI3K/Akt signaling pathway in the CDK1 expression and cell proliferation modulated by Tspan1 in human PDAC cells, the expression levels of PI3K, Akt and p-Akt were detected after Tspan1-overexpression, and Akt inhibitor MK-2206 2HCl were used to observe the protein expression of CDK1. The results showed that overexpression of Tspan1 could increase the content of Akt and p-Akt, so Tspan1-overexpression promoted the expression and phosphorylation of Akt, thus activate Akt in the BxPC3 cells and PNAC-1 cells. And, the suppression of Akt by MK-2206 2HCl significantly attenuated the increased CDK1 expression level induced by Tspan1-overexpression. In summary, these results indicate that Tspan1 modulated the expression of CDK1 via the activation of Akt.

Taken together, our study results indicated that Tspan1 promoted human PDAC cell proliferation by up-regulating CDK1 expression via activating Akt. The results indicate that Tspan1 may be used as an important biomarker for the diagnosis of PCC and prognosis. And Tspan1-silencing may serve as a potential therapeutic strategy to treat human pancreatic cancers.

## Availability of data and materials

The datasets used and/or analyzed during the current study are available from the corresponding author on reasonable request.

## Ethics Statement approval and consent to participate

The study was performed after obtaining approval from Cancer Hospital of China Medical University (Liaoning Cancer Hospital &Institute). Samples were taken from China Medical University with official written ethical consent.

## Acknowledgements

This research work is supported by the Liaoning Natural Science Foundation (2019-MS-211) and Scientific Research Project of Shenyang Science and Technology Bureau(18-014-4-75)

